# Biomimetic computer-to-brain communication restoring naturalistic touch sensations via peripheral nerve stimulation

**DOI:** 10.1101/2023.07.15.549130

**Authors:** Giacomo Valle, Natalija Katic Secerovic, Dominic Eggemann, Oleg Gorskii, Natalia Pavlova, Paul Cvancara, Thomas Stieglitz, Pavel Musienko, Marko Bumbasirevic, Stanisa Raspopovic

**Author notes:** Equal contribution (Giacomo Valle, Natalija Katic Secerovic).

## Abstract

Artificial communication with the brain through peripheral nerve stimulation recently showed promising results in people with sensorimotor deficits. However, these efforts fall short in delivering close-to-natural rich sensory experience, resulting in the necessity to propose novel venues for converting sensory information into neural stimulation patterns, which would possibly enable intuitive and natural sensations. To this aim, we designed and tested a biomimetic neurostimulation framework inspired by nature, able “to write” physiologically plausible information back into the residual healthy nervous system. Starting from the in-silico model of mechanoreceptors, we designed biomimetic policies of stimulation, emulating the activity of different afferent units. Then, we experimentally assessed these novel paradigms, alongside mechanical touch and commonly used, linear neuromodulations. We explored the somatosensory neuroaxis by stimulating the nerve while recording the neural responses at the dorsal root ganglion and spinal cord of decerebrated cats. Biomimetic stimulation resulted in a neural activity that travels consistently along the neuroaxis, producing the spatio-temporal neural dynamic more like the naturally evoked one. Finally, we then implemented these paradigms within the bionic device and tested it with patients. Biomimetic neurostimulations resulted in higher mobility and decreased mental effort compared to traditional approaches. The results of this neuroscience-driven technology inspired by the human body could be a model for the development of novel assistive neurotechnologies.

## Introduction

Loss of the communication between the brain and the rest of the body due to an injury or a neurological disease severely impacts sensorimotor abilities of disabled individuals. Often, they also experience the inability to sense their own body. The resulting low mobility and accompanying loss of independence cause a severe health problem and decline in quality of life with consequent necessary continuum of care. Recently developed neurotechnologies^1–3^ exploit direct electrical stimulation of the residual peripheral or central nervous system to restore some of the lost sensorimotor functions. Indeed, brain-computer interfaces (BCIs) exploiting implantable neural devices could potentially restore the bidirectional flow of information from and to the brain^1, 4, 5^. The implant of bio-compatible electrodes in the residual neural structures^6^, still functional after the injury, allows to create a direct communication channel. Indeed, neural stimulation of the peripheral somatic nerves (PNS)^7–10^, spinal cord^11–15^ or somatosensory cortex (S1)^16–18^ showed the ability to restore missing sensations, resulting in closed-loop neuroprostheses able to establish a bidirectional link between humans and machines. Sensory feedback restoration improved patients’ ability to use bionic limbs and increased its acceptance rate^5, 19–22^. However, the resulting dexterity of bionic hands is still far from that of natural hands in able-bodied individuals^23^, while mobility and endurance achieved with bionic legs are to be improved^24^. This is most probably due to multiple facts, among which that current neurotechnologies are falling short regarding the naturalness of induced sensations^25^, often resulting in unpleasant paresthesia. Indeed, common neuromodulation devices do not stimulate neurons based on the human natural touch coding or using model-based approach^26–28^, but rather with predefined constant stimulation frequency^29–31^. With these stimulation patterns, all elicited neurons are simultaneously activated, contrary to what happens with neural activity during in-vivo natural touch^32^. In fact, the natural asynchronous activation is driven in a part by the probabilistic nature of action potential generation in sensory organs, such as muscle spindles^33^ or touch afferents^34^, and in second part by the stochastic nature of synaptic transmission^35^. The synchronization, which generates an unnatural aggregate activity within the neural tissue, could be among the main reasons of perceived paresthesia percepts^8, 27, 36^. In fact, paresthetic sensations are likely to arise from this unnatural fibers activation^37^, and can be due to over-excitation of afferents or a cross-talk between them^38^. When caused by neuropathies, paresthesia is often chronic and do not improve over time, which might reflect an inability of central nervous system to learn how to interpret such aberrant neural responses^32^, making the use of electrical stimulation challenging. Moreover, it can interfere with the individual’s ability to sense and respond to other types of sensory information, such as touch or temperature. This can make it difficult to perform certain tasks or activities that require the use of multiple senses, or to interact with objects in the environment.

As a possible answer to this problem, the electrical stimulation built by mimicking the natural tactile signal (so called biomimetic sensory feedback^32, 39^) has shown to evoke more intuitive and natural sensations that better support interactions with objects, compared to usually used stimulation paradigms^40–42^. These biomimetic approaches might have the ability to electrically evoke aggregate population response similar to the natural one^23, 43^. Previous studies on natural touch suggests that somatosensory information about most tactile features is encoded synergistically by all afferent classes in the nerve^44^. Importantly, somatosensory cortex^45, 46^ (or even in cuneate nucleus^47^) are the earlies stages where signals coming from multiple fiber types converge and integrate with each other. It is allowing for the possibility that mimicking realistic neural responses of small mixed-type afferent populations will result in naturalistic patterns of cortical activation^43^, culminating in quasi-natural tactile percepts. However, despite the initial success of biomimetic approach in hand amputees where they outperformed classical non-biomimetic stimulation patterns, this approach was never tested in lower-limb amputees. Moreover, it was evaluated while performing tasks of daily living, or in more complex scenario than a single user with a single-channel stimulation. Furthermore, we are still lacking the understanding how these patterns are transmitted and interpreted in the first layers of information processing along the somatosensory neuroaxis.

To this aim, we develop a neuroprosthetic framework constituted by realistic in-silico modeling, pre- clinical animal validation and clinical testing in human patients with implants (**Fig.1**). Using this multifaceted approach, we are exploiting the architecture established by the development of validated model-based neurotechnology in human applications. More in detail, we first designed biomimetic neurostimulation strategies to restore somatosensory feedback by exploiting a realistic in-silico model of human afferents behavior (FootSim)^48^. This computational model can emulate the neural activity of the sensory afferents, innervating the plantar area of the human foot, in response to spatio-temporal skin deformations. It allowed designing neurostimulation patterns that potentially mimic relevant temporal features of the natural touch coding during walking. Together with the development of new stimulation paradigms, we assessed the major challenge: how specific artificial stimulation patterns translate into a neural signal and in how they travel along the somatosensory neuroaxis. With this goal, we stimulated the tibial nerves of decerebrated cats with cuff electrodes, while simultaneously recording neural activity (in Dorsal Root Ganglion (DRG) with a 32-channels Utah array and in spinal cord (L6) with a 32-channel shaft electrode). This setup allowed us to record and compare the electrically induced activity (in response to different patterns of nerve stimulation) with the response of neurons to mechanical touch. We validated this multifaceted approach through tests with three transfemoral amputees, with implants in tibial nerve. First, we compared the naturalness of the evoked artificial sensation using biomimetic and non-biomimetic encodings. Then, we implemented the biomimetic neurostimulation in a real-time, closed-loop neuroprosthetic leg, comparing its performance with respect to previously adopted neurostimulation strategies (linear and discrete neuromodulations). The patients performance were assessed during ecological motor tasks (i.e. a stairs walking task^49^ and a motor-cognitive dual task^50^).

**Fig. 1.**
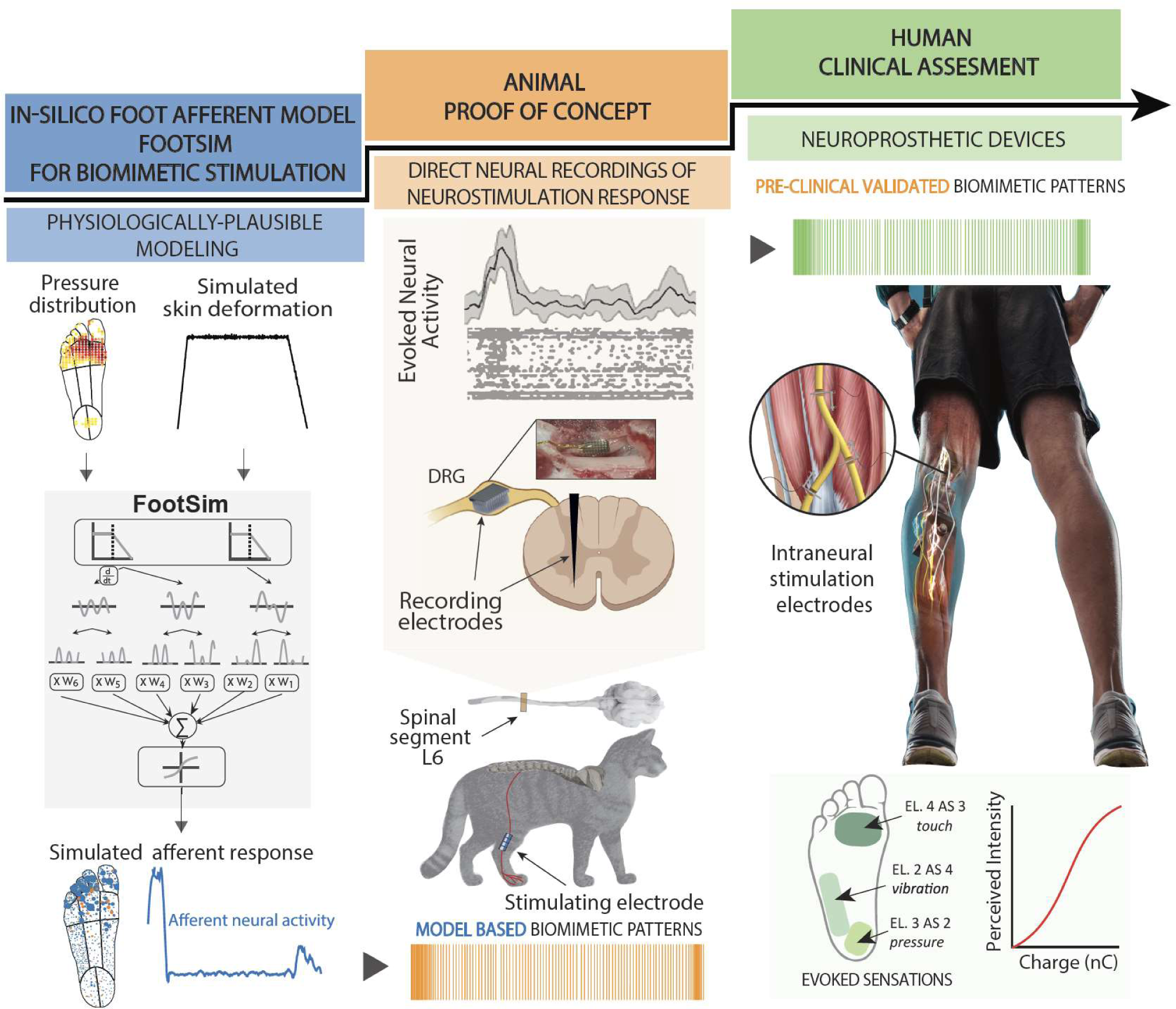
Neuroscience-driven development of a biomimetic neuroprosthetic device. The success development of a somatosensory neuroprosthesis is based on three main pillars: 1) In-silico models of the biological sensory processing have to be exploited for emulating the natural neural activation of the nervous system to external tactile stimuli (blue segment); 2) animal proof of concept allows for an experimental validation of the mechanisms behind the use of specific neurostimulation strategies defined with the use of modeling (orange segment); 3) A rigorous clinical validation of the biomimetic technology with implanted humans has to be performed in order to assess the functional outcomes in real-life scenarios (green segment). The results from the clinal trials will then allow to collect relevant data exploitable for improving computational modelling.

Both the animal and human experiments indicate that time-variant, biomimetic policies of artificial electrical stimulation should become the fundamental feature for design of next generation of neuroprostheses, able to directly communicate physiologically plausible sensations to the brain.

## Results

We exploited a trifold framework including modeling, animal and human experimentation (**Fig.1**) in order to design a neural bio-inspired stimulation strategy, effective for restoring somatosensation.

### Biomimetic neurostimulation paradigms are designed by exploiting a realistic in-silico model of foot sole afferents (FootSim)

We used the computational model of foot sole cutaneous afferents (FootSim)^48^ to design new biomimetic stimulation strategies.

FootSim is able to emulate the spatio-temporal dynamics of the natural touch considering the activation of all tactile afferents innervating the plantar area of the foot. This model is a plug-and- play tool, fitted on the human microneurography data, which models mechanical input from the external environment and gives on the output corresponding neural afferent activity (**Fig. 2a**). While setting up the environment, the user is populating the foot sole arbitrarily, depending on the case and envisioned usage. The foot can be filled with a realistic or modified distribution of a specific type of afferent, or alternatively, with the complete population. Different mechanical stimuli could be applied. We have the capability to simulate either a single mechanical stimulus applied to a specific position on the plantar side of the foot, or a scenario of a person walking. It can be achieved by extracting the pressure distribution across the entire foot sole at different time steps. (**Fig. 2a** left). The FootSim output can be structured in several forms. We can extract spike train of a single afferent, of summed population activity, or spatially represent the activity of the afferents placed in the foot sole by coding their firing rates with the area of the circle (**Fig. 2a** right).

**Fig. 2.**
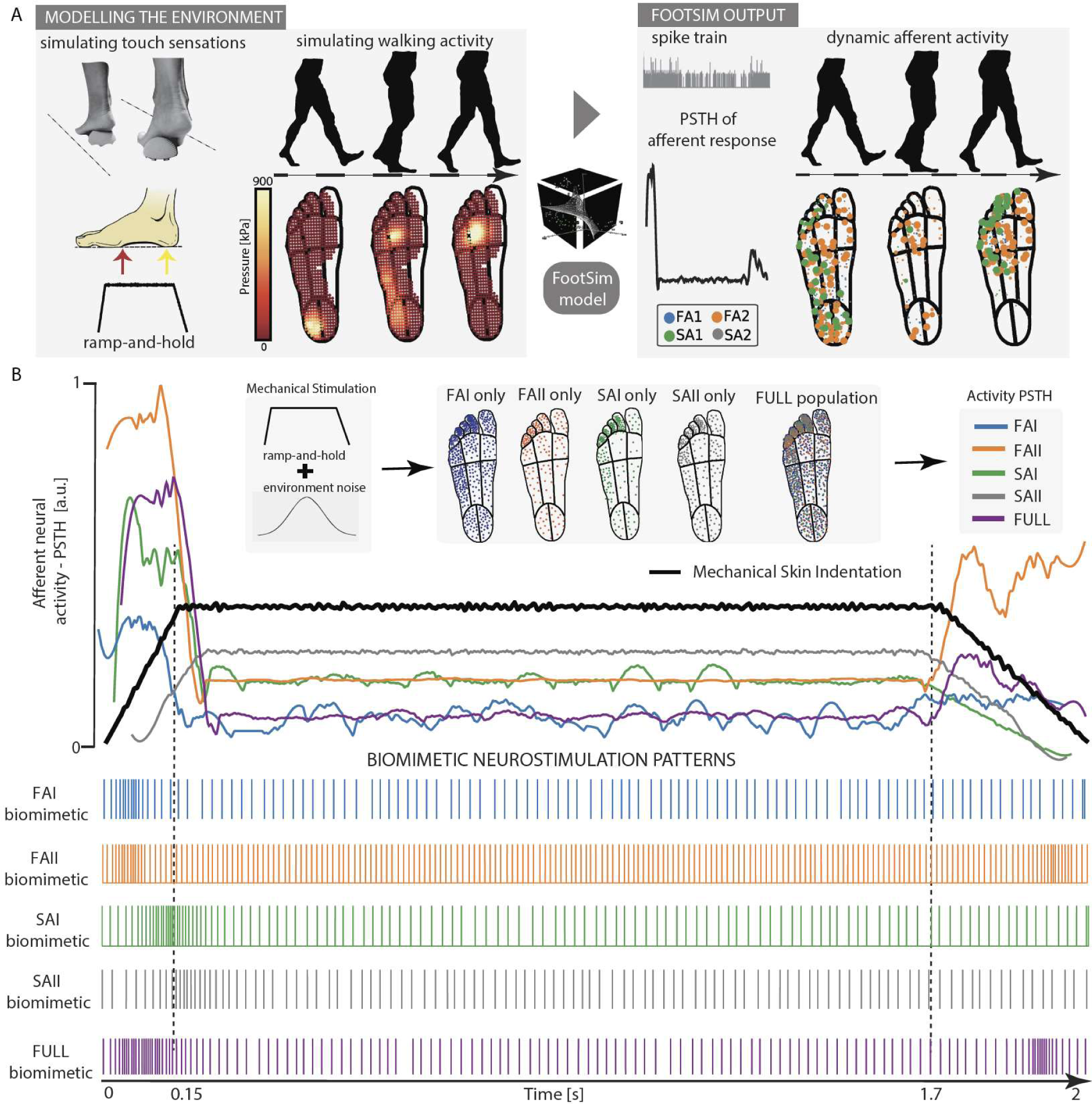
Biomimetic neurostimulation patterns designed using a realistic in-silico model of foot sole afferents (FootSim). (**a**) A schematic representation of practical use of FootSim plug-and-play model. Environment is modeled as a single, or continuous mechanical stimuli. The user can apply different types of stimulus, or simulate the walking scenario and the stimuli is given as an input to the model in form of pressure distribution across the foot sole. The output is afferent neural response that can be presented in several ways: as a single spike train, spatially represented on the foot sole by matching afferent firing rate with the area of the circle placed on the position of afferent, or as a populational response with peristimulus time histogram (PSTH). (**b**) Foot sole is populated with single type of afferent or with the whole realistic population (FAI/FAII/SAI/SAII/FULL population). We set the stimuli as a ramp-and-hold stimulus combined with the environmental noise and apply it on the whole foot area (black line). Neural responses of the whole applied population are given in the form of PSTH (colored lines). This was used as a function for the changes in frequency for defining biomimetic stimulating patterns. Amplitude remained constant in all biomimetic paradigms. All population distributions, afferent responses and respective biomimetic stimulation patterns are color coded: FAI: blue; FAII: orange; SAI: green; SAII: gray; FULL: purple.

When designing the biomimetic patterns, we also followed the aim to unveil if the naturalness can be coded in the neural responses specific to afferent type. We created 5 different scenarios by populating the foot sole with different types of afferents (**Fig. 2b**: FAI/FAII/SAI/SAII only), or with a complete population realistically existing in the human foot (**Fig. 2b**: FULL population). We applied a ramp-and-hold stimulus covering the whole foot sole with adding the environmental noise to mimic imperfection of the realistic pressure stimuli (**Fig. 2b** black line). We calculated the peristimulus time histogram (PSTH) merging all afferent responses based on the scenario (**Fig. 2b** colored lines). We used PSTH values to modulate the stimulating frequency, while keeping the amplitude constant, and create biomimetic neurostimulation paradigms. (**Fig. 2b**: FAI/FAII/SAI/SAII/FULL biomimetic).

### The neurostimulation dynamics is transferred through somatosensory neuroaxis

We recorded intra-spinal neural response signals and activity in dorsal root ganglion (DRG) in two cats to be able to compare bio- and non-bioinspired stimulation patterns and study their transmission through somatosensory axes. Cats were decerebrated for enabling the analysis of only reflex responses, avoiding the signal interference with voluntary movements^51^. Also, this procedure allows the testing without the use of anesthesia, that could potentially alter the neural responses. We implanted cuff electrode on tibial nerve for electrical stimulation and tuned the stimulation amplitude to be slightly above threshold. As multielectrode arrays showed to be the powerful tool for investigating the spinal cord processes^52^, we extracted neural signals from a dorso-ventral 32-channel linear probe implanted within the L6 spinal segment. Additionally, with UTAH array with 32 channels (**Fig. 3a**) we recorded neural signal in DRG at the L6 level.

**Fig. 3.**
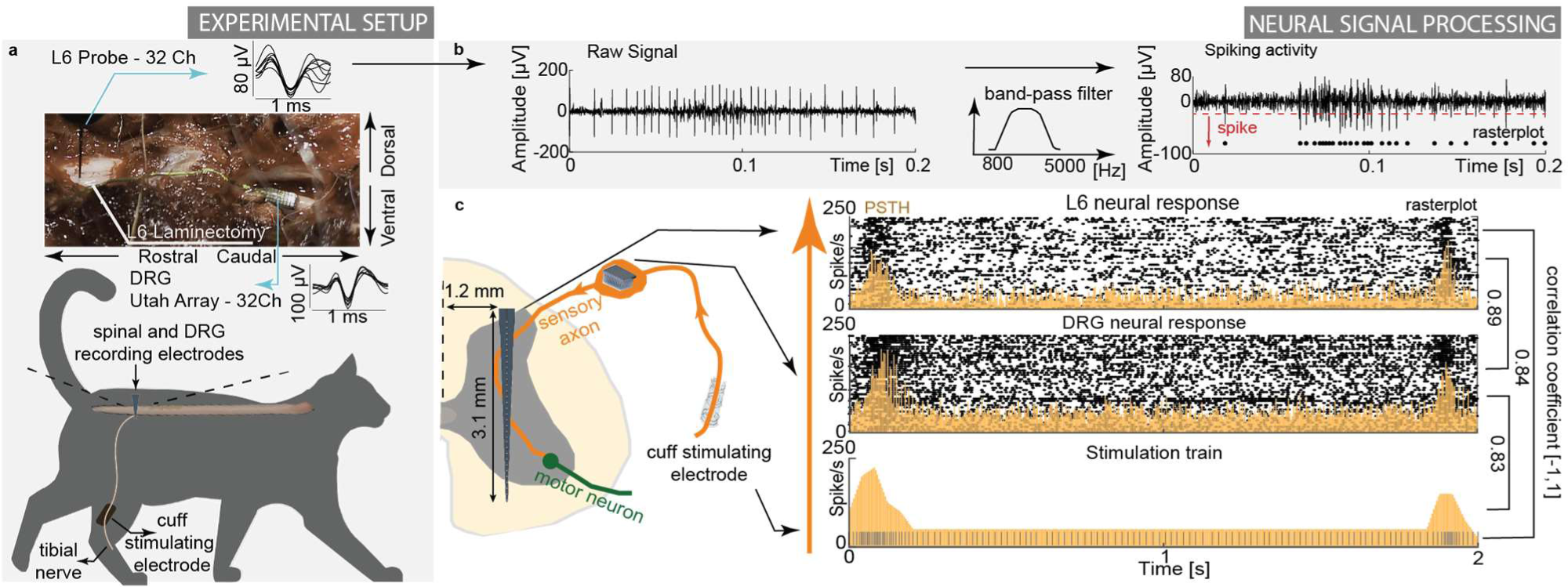
Purposefully designed experiments to study neural dynamics through neurostimulation. (**a**) Decerebrated cat experimental setup. We stimulated tibial nerve with cuff electrode and recorded neural response on the spinal level; upper part: exposed L6 vertebrae and dorsal root ganglion (DRG) with examples of recorded neural spikes from spinal linear electrode probe and DRG UTAH array. (**b**) Obtaining a multiunit neural activity. We filtered the signal to extract the spiking component and detect the neural action potentials using the thresholding algorithm. (**c**) Example of biomimetic stimulation paradigm and recorded response signal in one channel of spinal cord and DRG electrodes. Neural activity is presented and quantified with raster plot (black dots) and peri-stimulus time histogram (PSTH, yellow). Each row of the raster plot represents the response to a single biomimetic pattern (2s), while each dot corresponds to an action potential. Mean event rate (spikes/s) is defined as an average number of spikes within time frame of one bin (0.1 ms) across all single pulses of muscle nerve stimulation.

We tested the differences in neural dynamics that result from stimulating the tibial nerve with biomimetic paradigms and with tonic 50 Hz pattern that is commonly used in neuroprosthetics applications. We performed multi-unit threshold crossing analysis to identify the neural spiking activity (**Fig. 3b**), presented the results in form of rasterplot, and quantified them using peri-stimulus time histogram (PSTH) (see Methods).

The temporal dynamics of the neural activation pattern was highly correlated to the frequency of the neurostimulation train (**Fig. 3c**). In other words, by looking at the PSTH of single electrode channels, we can observe that multiple peripheral afferents responses followed the biomimetic pattern and thus encoded the artificial tactile information. Biomimetic pattern of activation was transmitted to the DRG maintaining the same spatio-temporal neural dynamics (R=0.83, p<0.05) and then also to the spinal cord (R=0.89, p<0.05). This evidence strongly supports the notion that electrical neural stimulation can serve as a highly efficient tool for generating artificial patterns of neural activations that can be effectively communicated to the upper regions of the somatosensory system. Indeed, biomimetic patterns of neurostimulation, induced at the peripheral nerve level, showed to evoke a very similar spatio-temporal neural dynamics in the spinal cord (R=0.84, p<0.05).

### Neural response evoked by biomimetic stimulation is more similar to the mechanically-induced activity than the one produced by tonic electrical stimulation

We base our hypotheses of evoking close to natural perception with biomimetic stimulation on the ability to code and replicate natural neural patterns. We recorded and compared the neural responses in DRG and spinal cord resulting from different types of electrical stimulation with the naturally induced neural activity, produced by touching the cat’s leg with the cotton bud.

Comparing the characteristics of the electrically-evoked neural dynamics resulting from applied biomimetic, non-biomimetic and natural stimulation confirmed previous theories^32, 39, 41^. Indeed, the temporal pattern of the evoked-response exploiting biomimetic neurostimulation encoding was more similar to the one generated by mechanical stimulation of the skin of the animal, than the one induced with tonic stimulation. We represented multi-unit spiking activity with PSTH (**Fig.4a**). We calculated mean neural activity produced during the period of electrical or natural stimuli for estimating the overall amount of information occupying the spinal cord and DRG. Natural touch and biomimetic stimulation resulted with similar values, while tonic stimulation inducted much higher activity in spinal cord and DRG. (**Fig.4a**, left). Shape of PSTH and its envelope gave an insight how neural activity is changing during the period of stimulation (natural, tonic or biomimetic). Biomimetic stimulation produces more similar activity as the natural touch compared to tonic stimulation (**Fig. 4a**, right). The encoded message is represented in the neural dynamics of activation. Results reveal that the information produced with biomimetic stimulation is matching better the natural touch neural coding then the commonly used tonic stimulation paradigm.

**Fig. 4.**
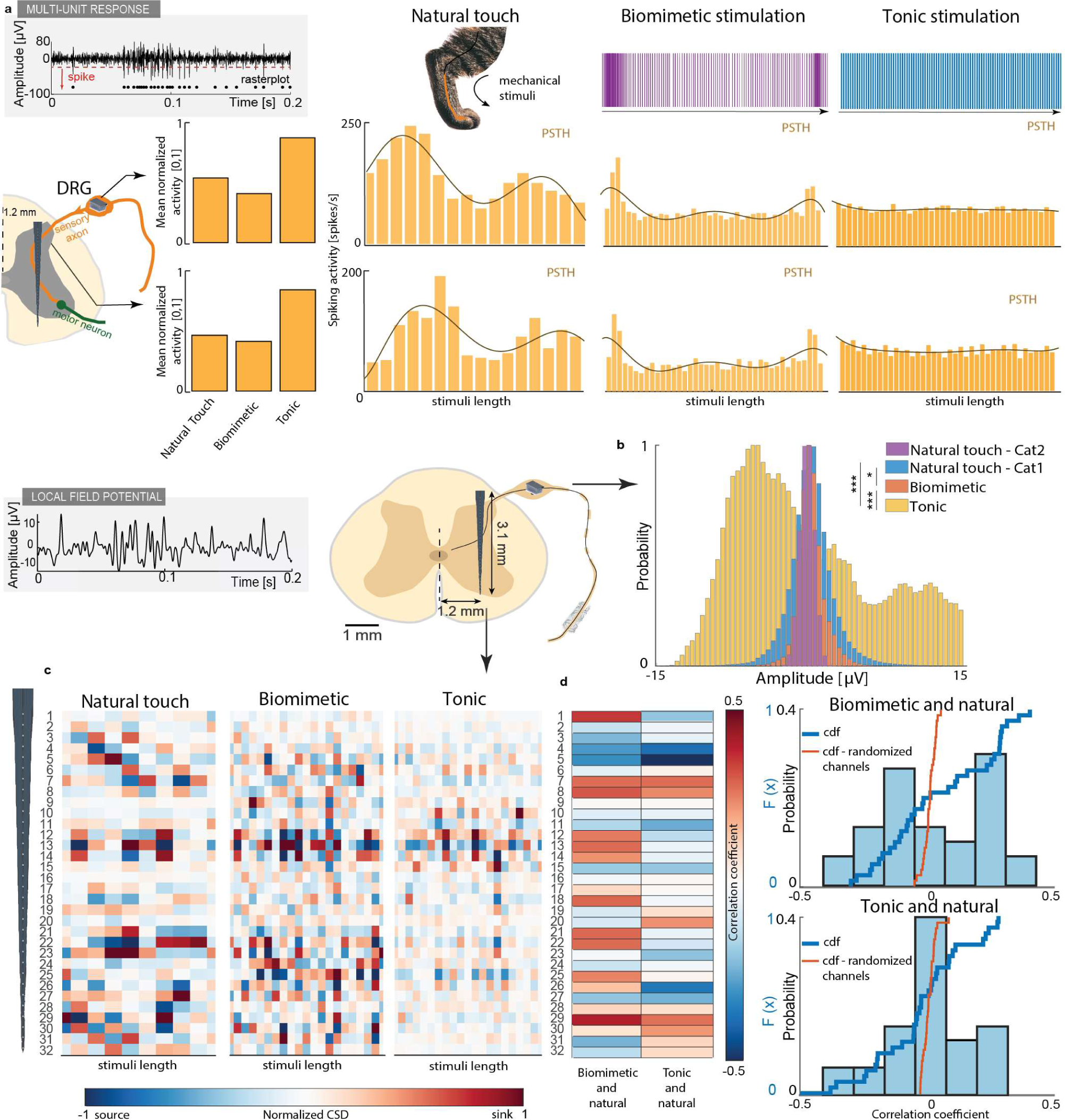
Neural response on biomimetic stimulation is more similar to the response to natural touch then to tonic stimulation. (**a**) Comparing multiunit neural activity as a response to natural touch, biomimetic or tonic (50 Hz) stimulation. We compared the signal recorded in DRG and spinal cord. Overall amount of neural activity during each condition is summed, normalized and presented with bars for comparison (left). Examples of spiking activity over time during each condition is presented using peri-stimulus time histogram (PSTH) (right) with the time bin of 50 ms. Brown lines represent the envelope of neural activity. (b) Comparing local field potential (LFP) recorded in DRG resulting from natural touch, biomimetic or tonic stimulation. Another natural touch response, recorded in cat 2, was added to analysis. We compared the distribution of specific signal amplitude values (*p<0.01; ***p<0.001). (c) Comparing current source density (CSD) calculated from LFP recorded in spinal cord resulting from natural touch, biomimetic or tonic stimulation. CSD is normalized for each condition and presented along the length of the electrode with 100 ms bin. (d) Left: correlation of CSD between biomimetic/tonic and natural touch condition, channel by channel, color coded. Right: Histogram and cumulative distribution function (cdf) of the correlation coefficient values resulting from comparing biomimetic/tonic stimulation and natural touch condition (top/bottom). Blue line represents the cdf when the recording channels are matched and compared. Red line corresponds to cdf when channels are randomly shuffled and compared.

Local field potential (LFP) reflects summed activity of small population of neurons represented by their extracellular potentials^53^ and they capture network dynamics^54, 55^. We performed the analysis of the trigger averaged LFP signal for different stimulating conditions. We extracted the DRG most active channels and investigated their amplitude variations. More in detail, we compared the amplitude distribution of recorded LFP (**Fig.4b**, see Methods). Tonic stimulation and natural touch responses showed statistically different amplitude distribution (p<0.001). The same conclusion arrived when we compared tonic and biomimetic stimulation (p<0.001). Response on biomimetic stimulation was more similar to the natural condition (p>0.05). As an addition, we tested natural touch condition in one more cat to investigate the cross-subject similarities of neural dynamics. The distributions of LFP amplitude were similar (p>0.05), showing that the naturally evoked response follows a specific, potentially generalizable trend, rather than being completely individual.

Current source density (CSD) is a technique for analyzing the extracellular current flow generated by the activity of neurons within a population of neurons. It can estimate the location and magnitude of current sources and sinks that contribute to the measured electrical signals. Therefore, we used it for comparing the spatial distribution of neural activity within a population of neurons along the array in the grey matter of spinal cord. We present the CSD estimated using local field potentials induced with biomimetic, tonic electrical stimulation, or natural touch (**Fig 4.c**). By looking at the spatial distribution of sinks and sources along the spinal axes, and comparing the overall resulting CSD, naturally induced touch response was more similar to the neural signal resulting from biomimetic stimulation (correlation coefficient 0.112) than to the one produced with constant, 50 Hz electrical stimulation (correlation coefficient 0.008). Additionally, we presented color coded channel-by-channel comparison of the resulting CSDs along the spinal electrode (**Fig 4d** left), and quantified the results with histogram and resulting cumulative distribution function (CDF) (**Fig 4d** right). CDF describes the probability that a random variable takes on a value less than or equal to a specified number. We used to compare distributions reflecting the comparison between CSD in different conditions. Tonic stimulation and natural touch produce neural responses with correlation coefficient very close to 0 in most of the channels, while that coefficient is higher for comparison between natural touch and biomimetic stimulation. In order to verify that this similarity is not produced by chance, we randomized the order of the channels in biomimetic and tonic electrical stimulation conditions and compared the recordings with the response of natural touch. It produced the correlation close to 0 for every electrode channel, confirming the validity of the used analyses.

Furthermore, we analyzed how much the neural signal is changing along the transversal spinal axes. We compared the correlation between the LFP in the first channel of intraspinal array and all the other channels (**Fig. S1**). In the natural touch condition, similarity between the neural activity is high in the first few channels and it is diminished when looking at more ventral recordings, in both animals. When nerve was electrically stimulated, similarity between neural activity recorded with the different channels through spinal array is high. The biomimetic neurostimulation elicited a less similarity along the spinal axes than tonic stimulation. Full population biomimetic pattern showed to be the more promising one compared to the paradigms created by mimicking response of specific afferent types. Despite being significant different from the natural touch, biomimetic stimulation based on aggregate population of afferent responses shares a striking similarity with it, setting it significantly apart from the tonic, 50 Hz stimulation.

### Biomimetic neurostimulation evokes more natural sensations than non-biomimetic neurostimulation paradigms

To test the functional implication of using biomimetic neurostimulations, we implemented and tested them in a human clinical trial. The scope was to firstly validate the biomimetic neurostimulation encoding assessing the quality of the evoked-sensations. Then, a real-time neuro-robotic device exploring biomimetic encoding strategies has to be compared to devices with previously-adopted encoding approaches in terms of functional performances. To this aim, three patients after transfemoral amputation (**Table S1**) were implanted with TIME electrodes in the tibial branch of the sciatic nerve (**Fig. 5a**). After a phase, called sensation characterization procedure, where all the 56 electrode active sites have been tested^36^, a subgroup of electrode channels were selected for this evaluation. Active sites eliciting sensations located in the frontal, central, lateral metatarsus and heel were identified (**Fig. 5b and Fig. S2**). In this way, the selected channels were electrically activating different groups of mixed afferents with projecting fields in different areas of the phantom foot (so with different distribution of innervating fibers).

**Fig. 5.**
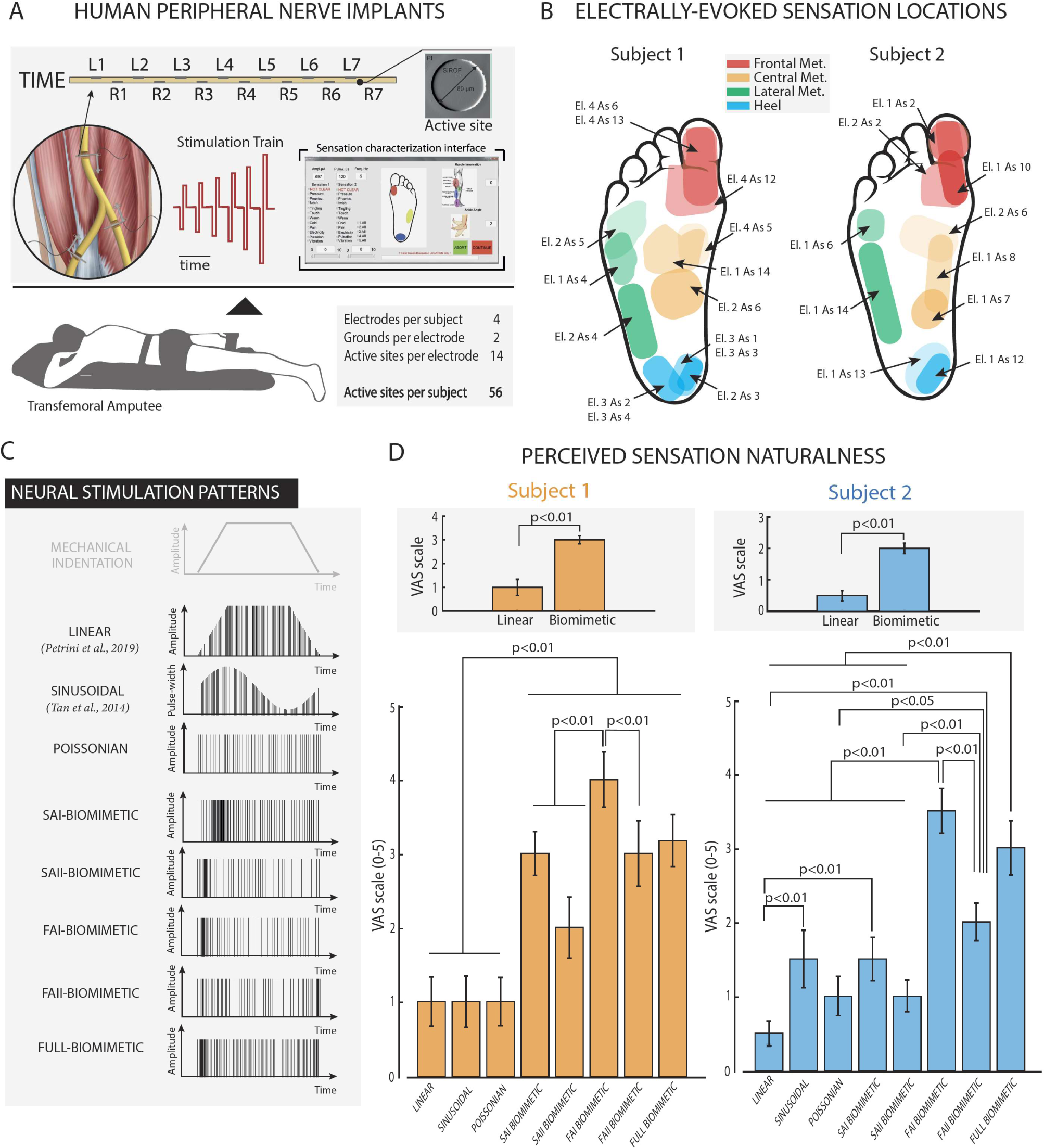
Biomimetic neurostimulations evoke more natural perceptions in implanted humans than non-biomimetic approaches. (**a**) Individuals with lower-limb amputation were implanted with TIME in their tibial nerves. The multichannel electrodes were used to directly stimulate the peripheral nerves evoking sensation directly referred to the phantom foot. (**b**) Projective fields map of two implanted subjects (1 & 2) related to the active sites adopted to electrically stimulate the nerves. Different colors show the 4 main regions of the phantom foot (Frontal, Lateral and Central Metatarsus, and Heel). (**c**) Biomimetic and non-biomimetic neurostimulation strategies adopted for encoding a mechanical indentation of the foot sole. Linear neurostimulation is taken from ^36^ and Sinusoidal neurostimulation by ^10^ (**d**) Naturalness ratings (VAS scale 0-5) of the perceived sensation elicited exploiting different stimulation strategies in two subjects. Insets: Group comparison between linear vs biomimetic stimulations.

Then, multiple strategies, encoding a mechanical skin indentation, have been adopted to deliver neurostimulation trains through each selected channel of the intraneural implants (**Fig.5c**). The participants were asked to report the perceived sensation naturalness using a visual analogue scale (VAS) between 0 (totally non-natural sensation) and 5 (totally natural sensation – skin indentation) ^41, 56^. In all the three implanted subjects and considering all the active sites tested (with different projected fields), the biomimetic neurostimulation patterns elicited sensations more natural than the linear neurostimulation encoding (3±0.18 with Biomimetic compared to 1±0.35 in Linear for S1, 2±0.16 with Biomimetic compared to 0.5±0.17 in linear for S2, and 2±0.36 with Biomimetic compared to 1±0.18 in linear for S3 across all electrode tested, p<0.01) (**Fig.5d and Fig.S2**) that was previously adopted in multiple neuroprosthetic applications^8, 27, 36^. Moreover, biomimetic-based encodings often resulted in more natural perceived sensations compared to both sinusoidal (pulse width-variant) and Poisson (frequency-variant) neurostimulation strategies (p<0.05), indicating the importance of inducing a neural activation dynamic mimicking the natural biological code.

Notably, although multiple biomimetic-like paradigms have been tested (SAI-, SAII-, FAI-, FAII-like and Full biomimetic), none of them proved to be better. Although biomimetic stimulation was always eliciting more natural sensations than one parameter adopted encoding, analyzing the results per location in both subjects (**Fig.S3**) did not show any clear evidence of an optimal biomimetic encoding schema. This was probably caused by the different composition of the fibers activated by the electrode channels in the different foot regions^57^. In fact, the perceived areas were different according to the active site selected to stimulate, indicating a different group of mixed afferents recruited by the neurostimulation. We hypothesized that not only the proportion of SA and FA fibers is relevant, but also their role in encoding touch information in that specific region.

These findings highlighted how biomimicry is a fundamental feature of the electrical neural stimulation for successfully restoring more natural somatosensory information.

### Biomimetic neurostimulation on a neuro-robotic device allows for a higher mobility and a reduced mental workload

Aiming to develop a neuroprosthetic device able to replace the sensory-motor functions of a natural limb as much as possible, this biomimetic neurostimulation was then implemented in a real-time robotic system. This wearable system was composed by: i) a sensorized insole with multiple pressure sensors; ii) a microprocessor-based prosthetic knee with a compliant foot (Ossur, Iceland); iii) a portable microcontroller programmed with the biomimetic sensory encoding algorithms; iv) a multichannel neurostimulator; v) intraneural electrodes implanted in the peripheral nerves (TIMEs).

The neuroprosthetic device was working in real-time being able to record pressure information from the wearable sensors, while the patient was walking, and converting them in patterns of biomimetic neurostimulation delivered through the TIMEs (see Method for implementation details). In this way, the users were able to perceive natural somatotopic sensations coming directly from the prosthetic leg without any perceivable delay.

After the implementation, we assessed the effects of exploiting the biomimetic encoding (BIOM) in a neuro-robotic device compared to a linear (LIM) or a time-discrete (DISC) neurostimulation strategy. In the LIN, the sensors’ readouts were converted in neurostimulation trains following a linear relationship between applied pressure and injected charge^27, 36^. In case of DISC, short-lasting, low-intensity electrical stimulation trains were delivered synchronously with gait-phase transitions ^58, 59^. Also the condition without the use of any neural feedback (NF) was included in the motor paradigms as a control condition.

The neuroprosthetic users were thus asked to perform two ecological motor tasks: Stairs Task (ST)^36, 49^ and Cognitive Double Task (CDT)^50^.

In ST, results indicated that, when exploiting biomimetic neurostimulation in a neuro-robotic leg, both users improved their walking speed (4.9±0.1 for S1 and 4.3±0.4 for S2 laps/session) compared to LIN (4.5±0.1, p<0.05 for S1 and3.8±0.1, p<0.05 for S2 laps/session), DISC (4.6±0.1, p<0.05 for S1 and 3.6±0.1, p<0.05 for S2 laps/session) and NF (4.3±0.1, p<0.05 for S1 and 3.5±0.1, p<0.05 for S2 laps/session) conditions (**Fig.6a**). Interestingly, also the self-reported confidence (VAS scale 0-10) in walking on stairs was increased, when the participants were exploiting the neuroprosthetic device with biomimetic neurofeedback (9.75±0.26 for S1 and 6±0.3 for S2) compared to LIN (8.75±0.62, p<0.05 for S1 and 5.37±0.23, p<0.05 for S2), DISC (7.83±0.39, p<0.05 for S1 and 5.17±0.25, p<0.05 for S2) and NF (6.67±0.49, p<0.05 for S1 and 3.83±0.25, p<0.05 for S2) conditions (**Fig.6a**).

**Fig. 6.**
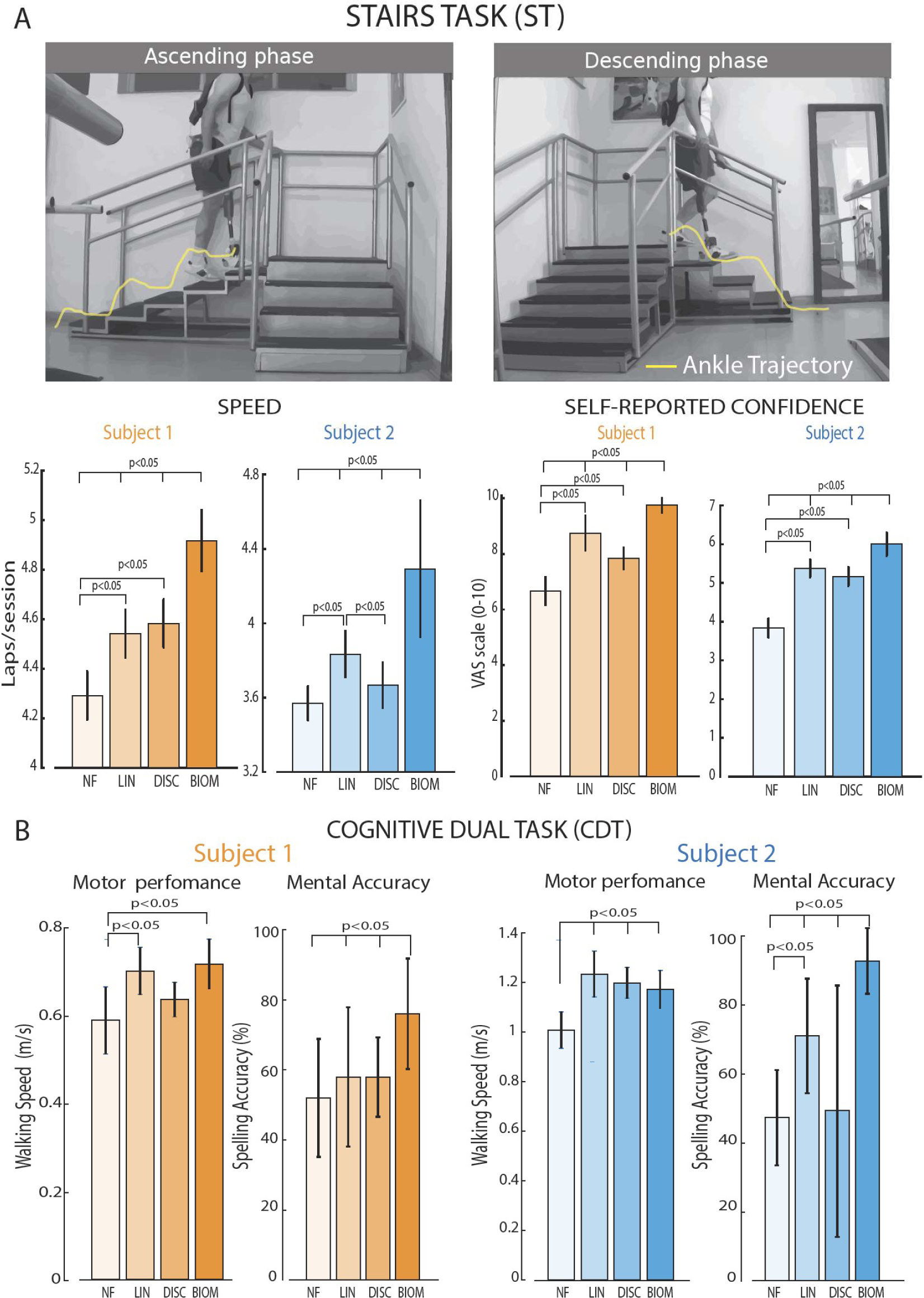
Real-time biomimetic neural feedback allows for higher speed and lower cognitive workload while walking. (**a**) In the Stairs Task (ST), the subjects (1 & 2) were asked to walk over stairs in both ascending and descending directions. (**b**) Speed (Laps/session) and self-reported confidence (VAS Scale 0-10) were measured in ST. (**c**) Motor performance (Walking Speed – m/s) and Mental Accuracy (Spelling Accuracy - %) of Subject 1 & 2 in the Cognitive Dual Task (CDT). In both tasks, conditions are NF (No Feedback), LIN (Linear Neurostimulation), DISC (Discrete Neurostimulation) and BIOM (Biomimetic Neurostimulation).

In the CDT, both participants showed a higher mental accuracy in BIOM compared to the other conditions (p<0.05 in both subjects), while maintaining the same walking speed. In particular, the mental accuracy of S1 was 76±16% in BIOM, 58±20%in LIN, 58±11% in DISC and 52±17% in NF while in S2 94±9.6% in BIOM, 72±17% in LIN, 50±37% in DISC and 48±14% in NF. Notably, the walking speed was always higher in the feedback conditions compared to NF for S2 (p<0.05) and in BIOM and in LIN for S1 (p<0.05). As expected, without adding a secondary task, no difference was observed in the walking speed among the conditions in both subjects (p>0.1, **Fig.S4**). These findings indicated a higher decrease in mental workload, while the users were performing two tasks simultaneously (one motor and one cognitive) in the moment that a more bio-inspired neural stimulation was exploited in a neuro-robotic device.

## Discussion

### Multi-level approach for designing new stimulation strategies that would minimize paresthesia sensations

In this study, we designed, developed and tested a neuro-robotic device exploiting model-based biomimetic neurostimulations in people with limb amputation. Due to a multilevel framework, it was possible to design and test effective bio-inspired neurostimulation paradigms to elicit more natural feelings and better understand the reasoning behind the use of biomimetic approaches in the neuroprosthetic field. Indeed, thanks to realistic in-silico modeling of the foot touch coding, precise neural stimulation patterns were defined that could accurately emulate the firing of the cutaneous mechanoreceptors. Single-fiber (SAI, SAII, FAI, FAII) and mixed-fibers (FULL) type patterns have been implemented to encode a mechanical skin indentation into a neural stimulation policies. We modulated the stimulation frequency based on the fiber dynamics of activation since it showed to be beneficial for shaping the artificial touch for bionic limbs^60^.

### Comparing neural responses induced with natural touch and electrical nerve stimulation

The purposely-designed animal experiments allowed us to compare the neural dynamics as a response to natural touch, biomimetic or tonic electrical stimulation. The recordings in decerebrated cats via multiple neural interfaces along their somatosensory neuroaxis (somatic nerve, DRG and spinal cord) showed that biomimetic neurostimulations evoked spatio-temporal characteristics of the afferents’ response more similar to the naturally induced one than tonic stimulation. These biomimetic patterns are going towards avoiding highly synchronized activity in spinal circuits that could saturate them and limit the possibility to perceive touch sensations restored with electrical stimulation^61^. This is clear evidence of the effect of bio-inspired stimulation dynamics on the neural afferents activation, showing the possibility to artificially encode natural sensory messages into the nervous system. Indeed, previous researches have hypothesized the adoption of complex spatiotemporal patterns mimicking natural peripheral afferents activity^32, 39^. This approach was also proposed for cortical activity modulation using intracortical microstimulation (ICMS) to convey feedback of touch^42, 62^ or of the entire movement trajectory (natural proprioceptive sensation)^63^. Likewise, they also assumed that exploiting an ICMS interface that mimics natural sensations would be faster, and ultimately more effective than learning arbitrary associations with unnatural sensations or arbitrarily modulated ICMS^64^. Our study validates these hypotheses on the use of biomimetic encoding in PNS neuroprostheses. However, our experimental setup was focused to understanding the first layer of processing information coming from the periphery, until the spinal cord. Going higher in this direction, cortical responses (LFPs) to biomimetic peripheral nerve stimulation using interfascicular electrodes have been recently measured in a monkey^65^. The authors showed that constant frequency stimulation produced continual phase locking, whereas biomimetic stimulation produced gamma enhancement throughout the stimulus, phase locked only at the onset and release of the stimulus. This cortical response has been described as an “Appropriate Response in the gamma band” (ARγ). Regarding the sensory restoration in bionics, multichannel biomimetic ICMS showed to provide high-resolution force feedback^42^ and more localizable sensations^66^ in implanted humans. We believe future experimental work should extend these findings investigating neural processes caused by electrical stimulation in gracilis nucleus (or cuneate for the upper limb), thalamus or in somatosensory cortex, in particular in humans.

### Biomimetic stimulation in neuromodulation devices is beneficial both for the perceived sensation and its functionality

These biomimetic neurostimulation strategies were tested in three human subjects with transfemoral amputation implanted in their leg nerves with intraneural electrodes. All the participants reported to feel more natural sensations, when stimulated with biomimetic encodings with respect to standard neuromodulation patterns from every stimulation channel on the electrodes. Neural stimulation gradually recruit all the sensory afferents within the fascicle^67, 68^ depending on both distance from the electrode (threshold proportional to square of distance) and their diameter (threshold proportional to 1/square root of fiber diameter)^69^. Therefore, each stimulation pulse delivered through the active site is likely to recruit a mix of sensory afferents types, even if clustered^70^. For this reason, how many and what tactile afferents will be stimulated by a given stimulation pattern through a specific electrode is unknown a priori. This might be the reason why different types of biomimetic encoding were reported as more natural by the participants according to the perceived foot location (**Fig.S3**) and, therefore, to the clusters of recruited afferents. This phenomenon can also reveal the typology of sensation reported, while specific types of afferents were activated by neurostimulation^38, 71^ (flutter, vibration, touch). Similarly, it could explain why with simpler encoding (at the threshold level) the electrically-evoked sensation can be sometimes reported as natural^8, 10, 36, 72^.

Finally, we implemented these algorithms in robotic prosthetic devices in a real-time fashion comparing their functional performance with previously-proposed technologies. Biomimetic neuroprosthetic legs allowed for a faster stair walking and a decreased mental workload in a double task paradigm in both subjects. These findings demonstrated that biomimetic encoding is relevant for device functionality and thus to enhance the beneficial effect of this intervention. In particular, a significant boost in mobility, on a difficult everyday life task as the stairs, is very relevant for people with lower-limb amputation. This improvement is likely connected to reported higher confidence in the prosthetic leg with biomimetic sensory feedback^49^. The amputee is able to instantly sense the position of his leg with regard to the ground more naturally, which allows him to transition faster from heel strike to loading his prosthetic leg^73^. Confidence and mobility have been previously proposed to be among the clearest and simplest parameters showing the impact of sensory feedback on gait^49^. Regarding the CDT, it represented a real-life scenario of multiple simultaneous tasks. It allowed us to obtain an objective measure of the better cognitive integration of the prosthesis with biomimetic neurostimulation^22, 50^, since both amputees improved their mental accuracy. In addition to our results, previous studies have also preliminarily shown these improvements in manual dexterity and object recognition in upper-limb amputees exploiting robotic hand prostheses^40, 41^.

### Limitations of the study

Biomimetic stimulation paradigms are developed based on FootSim model output, where we simulated applying ramp-and-hold mechanical stimuli on the sole of the foot. The natural mechanical stimulation provided in the experimental protocol with decerebrated cats do not exactly replicate the same stimuli. However, the complexity and length of the surgery and experiments, were extremely high, so we applied the stimuli suitable for performing in a reasonable time frame. To do so we delivered rapid tactile stimuli to the cat leg with a cotton swab, as in this way we are able to superficially activate multiple sensory fibers in tibial nerve, increasing therefore the likelihood to effectively elicit the signals that we can effectively record. Instead, touching spatially distinctive foot sole areas, could result in multiple positions not eliciting any activity that we could record, therefore exceeding the available experimental time. Future tests should examine the comparison of PNS with different types of natural tactile stimulation.

Relevant limitations regarding the FootSim model are connected to the fact that the model is not incorporating shear forces or lateral sliding but simulates them as a quasi-continuous stress. This simplification implies reduced accuracy of predicting the SA2 type afferents’ responses, which transmit the information about skin stretch.

Finally, even though we tested the biomimetic patterns in the closed-loop neuroprosthetic system, these paradigms were based on the model outputs run offline, while we believe that the stimulation strategies should be defined in the real time from the output of the model. Engineering efforts are needed to make this neuroprosthetic system fully-functionable in real time and, in the future, also fully implantable.

### Use of this framework in future biomimetic neurostimulation devices

Presented neuromodulation framework based on biomimetic encoding could also be very relevant for other neuroprostheses in the CNS (e.g., Deep Brain Stimulation^74^, epidural stimulation^13^, ICMS^5, 18^) and for bioelectronic medicine applications (e.g. vagus stimulation^75^, stimulation of the autonomic nervous system^76^) having the same necessity to evoke a natural pattern of activation in a certain nervous district using artificial electrical stimulation. Indeed, the biomimetic approach has been proven to be effective for improving functional performance in other type of neural prostheses (e.g., enhanced speech intelligibility for cochlear implants^77^; improved restoration of gaze stability in vestibular prostheses^78^). We believe that approach based on the in-silico modelling of the desired neurological function, followed by animal validation prior to human testing evaluating both the perceived quality of sensations and performance while doing daily tasks, will become the standard framework for the development of the novel neuroprostheses.

Considering future biomimetic neuro-robotic devices restoring fully-natural sensations, spatial patterning can be achieved by stimulating different electrodes with spatially displaced projection fields, while temporal patterns can be elicited by temporally modulating the stimulation parameters delivered through each electrode, as proposed in our study. However, the extent to which artificially evoked neural activity must mimic that of the natural afferent inputs in order to be fully-exploitable also for more complex tactile features^26, 79^ (textures, object stiffness, shape, etc.) or proprioception remains a critical question.

Here we evaluated multiple types of biomimetic patterns that were developed using the distinct response characteristics of individual afferent types. When we stimulated the entire nerve during animal experiments, the biomimetic pattern based on the aggregate afferent response (FULL biomimetic) showed to be the most promising one compared to its natural counterpart. Notably, when these paradigms were delivered using intraneural electrode in humans, smaller clusters of mixed afferents have been selectively activated by the different channels. Interestingly, the naturalness of the sensation, for the same encoding strategies, changed accordingly to specific areas of the foot sole. It suggests that the imposition of the aggregate dynamics for inducing natural sensations is not optimal for every fiber cluster recruited. It seems to depend on the distribution of activated afferents (mechanoreceptors) and their specific role in the sensory processing. We believe that neurostimulation strategies should be informed by computational modelling emulating realistic dynamic conditions.

In conclusion, our collected evidence not only amplifies the remarkable impact of biomimetic signal encoding from a scientific perspective, but it also holds immense promise in heralding the advent of the next generation of neuroprosthetic devices. New technologies, inspired by nature, have a potential to fully emulate natural neural functions lost after a disease or an injury. The possibility to naturally communicate with the brain will open new doors for science in multiple fields.

## Methods

### Modeling of all tactile afferents innervating the glabrous skin of the foot (FootSim)

In this study, we used FootSim^48^, in-silico model of the afferents innervating the foot sole that simulates the neural responses on arbitrary mechanical stimuli. It is composed of the two parts i) mechanical part, calculating the deformation of the skin by applied stimulus and converting it into the skin stress ii) firing models that generate spiking output for individual fibers of different afferent classes. Each firing model contains 11 unique parameters. The model is fitted on a dataset of tactile afferents exposed to a wide range of vibrotactile stimuli at different frequencies and amplitudes, recorded in humans using microneurography. We fitted several models for each afferent type, reflecting partially the natural response variability of different afferents observed in the empirical data.

### Design biomimetic neural stimulations using FootSim

We designed 5 types of biomimetic patterns, based on the cumulative responses of specific afferent types. In FootSim model, we populated the foot sole with only one type of afferents (FA1/FA2/SA1/SA2) or with a complete population of afferents (FULL biomimetic), following their realistic distribution. We applied 2 s stimuli covering the whole area of the foot. We combined ramp-and-hold stimuli (0.15 s on phase, 0.3 s off phase) with low-amplitude environmental noise (up to 0.5% of maximum amplitude of ramp-and-hold stimuli). FootSim model estimated the response of each single afferent placed on the sole of the foot. We aggregated the spiking activity of all units, smoothed the obtained function over time and used it as a modulated frequency of stimulation for each biomimetic pattern. Amplitude and pulse-width of stimulation, identified during the electrode mapping procedure, were kept constant along the train.

### Animal surgical procedure

Experiments were carried out on 2 adult cats of either sex (weighing 2.5-4.0 kg). All procedures were conducted in accordance with protocols approved by the Animal Care Committee of the Pavlov Institute of Physiology, St. Petersburg, Russia, and adhered to the European Community Council Directive (2010/63EU). The surgical procedures were similar to those in our previous studies^80, 81^. The cats were deeply anesthetized with isoflurane (2-4%) delivered in O_2_. For the induction of anesthesia, xylazine (0.5 mg/kg, i.m.) was injected. The level of anesthesia was monitored based on applying pressure to a paw (to detect limb withdrawal), as well as by checking the size and reactivity of the pupils. The trachea was cannulated, and the carotid arteries were ligated. The animals were decerebrated at the precollicular-postmammillary level to assure the pure sensory recordings, without influence of the higher structures. An access to tibial nerve, laminectomy in corresponding segments for intraspinal and DRG recording of neurons were performed (Fig. 3). Cuff electrode (Microprobes for Life Science, Gaithersburg, MD 20879, U.S.A) is placed after the careful dissection from surrounding tissues, around the common trunk of tibial nerve. The exposed dorsal surface of the spinal cord was covered with warm paraffin oil. Linear shaft electrodes with 32 channels (Neuronexus, Ann Arbor, MI, U.S.A.) are carefully implanted at the spinal level L6, using stereotaxic frame. DRG implant has been performed by implanting the 32-channel UTAH array (Blackrock Microsystems, Salt Lake City, UT, U.S.A.), though the pneumatic injection pistol. Anesthesia was discontinued after the surgical procedures that lasted up to 20 hours. The experiments were started 2–3 h thereafter. During the experiment, the rectal temperature and mean blood pressure of the animals were continuously monitored and kept at 37 ± 0.5°C and above 80 mmHg.

### Electrophysiology in decerebrated cats

Through the contact sites of the cuff electrodes, we delivered single pulses of cathodic, charge balanced, symmetric square pulses (with pulse width of 0.5 ms). We provided the stimulation using AM stimulators Model 2100 (A-M Systems, Sequim, WA, USA). Electromyographic and neural signals were acquired using the LTR-EU-16 recording system with LTR11 ADC (L-Card, Moscow, Russia) and the RHS recording system with 32-channel headstages (Intan Technologies, Los Angeles, CA, U.S.A.) at a sampling frequency of 25 and 30 kHz respectively. We tuned the stimulation amplitude by observing the emergence of clear sensory volleys in the dorsal spinal cord in response to low-frequency stimulation.

We applied 5 types of biomimetic stimulation paradigms, repeating every pattern 90 times. Natural touch condition was applied by rubbing the cat’s leg with cotton swab and was repeated 5 times.

### Analysis of the animal neural data

After acquiring animal neural data, we applied all detailed analysis offline, as following:

#### Pre-processing

We filtered raw signals recorded with 32-electrode array implanted in the spinal cord, as well as signals documented with 32-channel Utah array in dorsal root ganglion with comb filter to remove artefacts on 50 Hz and its harmonics. We designed a digital infinite impulse response filter as a group of notch filters that are evenly spaced at exactly 50 Hz. We removed signal drift with a high-pass 3^rd^ order Butterworth filter with a 30 Hz cutoff frequency. High amplitude artifacts were detected when the signal crossed a threshold equal to 15σ, where we estimated background noise standard deviation^82^ as σ = median |x| 0.6745. Detected artifacts were zero-padded for 10 ms before and after the threshold crossing. We extracted neural signal of 2 s recorded during stimulation with every defined paradigm. Natural touch condition produced response of 1 s and the signal where neural activity was observable was extracted.

#### Identification of local field potential

We isolated local field potentials by band passing the neural signal between 30 Hz-300 Hz and averaged the signal over multiple stimuli pattern repetitions.

#### Characterization and quantification of neural spiking activity

We extracted neural spiking activity by applying a 3rd order Butterworth digital filter to the raw signal, separating the signal in frequency range from 800 Hz to 5000 Hz. We detected the spikes using unsupervised algorithm^83^. We determined the threshold value separately for each recording channel. To detect the accurate threshold value, we concatenated all data sets recorded in one place (spinal cord/DRG) that we aim to analyze in a single file. All analyzed data sets were concatenated in a single file in order to detect proper threshold values. Threshold for detection of action potentials was set to negative 3σ for signals recorded in the spinal cord and 4σ for signals recorded in the DRG, where σ = median |x| 0.6745 which represents an estimation of the background standard deviation.

Multiunit activity is presented in form of rasterplot and quantified with peri-stimulus time histogram (PSTH). Each dot in rasterplot represents a single detected spike. Every rasteplot row corresponds to the intra-spinal or intra-cortical activity perturbed with a single muscle nerve stimulus pulse. PSTH is quantified with mean event rate, defined as the average number of spikes across all single pulses of muscle nerve stimulation, within defined time frame.

### Patient recruitment and surgical procedure in humans

Three unilateral transfemoral amputees were included in the study. All of them were active users of passive prosthetic devices (Ottobock 3R80) (**Table S1)**. Ethical approval was obtained from the institutional ethics committees of the Clinical Center of Serbia, Belgrade, Serbia, where the surgery was performed (ClinicalTrials.gov identifier NCT03350061). All the subjects read and signed the informed consent. During the entire duration of our study, all experiments were conducted in accordance with relevant EU guidelines and regulations.

Four TIMEs^84^ (14 active sites each) were obliquely implanted in the tibial branch of the sciatic nerve of each subject. The surgical approach used to implant TIMEs has been extensively reported elsewhere^7^. Briefly, under general anesthesia, through a skin incision over the sulcus between the biceps femoris and semitendinosus muscles, the tibial nerve was exposed to implant 4 TIMEs. A segment of the microelectrodes cables was drawn through 4 small skin incisions 3 to 5 cm higher than the pelvis ilium. The cable segments were externalized (and secured with sutures) to be available for the transcutaneous connection with a neural stimulator. After 90 days, the microelectrodes were removed under an operating microscope in accordance with the protocol and the obtained permissions.

This study was performed within a larger set of experimental protocols aiming at assessing the impact of the restoration of sensory feedback via neural implants in leg amputees during a 3-month clinical trial^7, 36, 49, 50, 85^. The data reported in this manuscript was obtained in multiple days during the 3-months trial in three leg amputees.

### Intraneural stimulation for evoking artificial sensations

Each of the TIMEs (latest generation TIME-4H) implanted in the three amputees was constituted by 14 active sites (AS) and two ground-electrodes. Details concerning design and fabrication can be found in ^86, 87^. For each subject, 56 electrode channels were then accessible for stimulation on the tibial nerve. During the characterization procedure the stimulation parameters (i.e. amplitude and pulse-width of the stimulation train), for each electrode and AS, were recorded. The electrodes were connected to an external multichannel controllable neurostimulator, the STIMEP (Axonic, and University of Montpellier) ^88^. The scope of this procedure was to determine the relationships between stimulation parameters and the quality, location, and intensity of the electrically-evoked sensation, as described by Petrini et al. ^36^. In brief, the injected charge was linearly increased at a fixed frequency (50 Hz ^36^) and pulse-width by modulating the amplitude of the stimulation for each electrode channel. In case the stimulation range was too small for the chosen pulse-width and the maximum injectable current, the pulse-width was increased, and the same procedure was repeated. When the subject reported to perceive any electrically-evoked sensation, the minimum charge (i.e. perceptual threshold) was registered. The maximum charge was collected in order to avoid that the sensation became painful or uncomfortable for the subject. This was repeated five times per channel and then averaged. Perceptual threshold and maximum charge were obtained for every electrode channel and have been used to choose the stimulation range. For each AS, the maximum injected charge was always below the TIME’s chemical safety limit of 120 nC ^89^. All the data were collected using a custom-designed psychometric platform for neuroprosthetic applications. It indeed allows to collect data using standardized assessment questionnaires and scales, and to perform measurements over time. The psychometric platform is user-friendly and provides clinicians with all the information needed to assess the sensory feedback ^90^.

### Assessment of sensation naturalness

We first characterized the subjects’ rating of the perceived naturalness of the stimulation delivered through TIMEs in S1, S and S3. We injected biphasic trains of current pulses lasting 2 s with an increasing phase (0.5 s), a static phase (1 s) and a decreasing phase (0.5 s) via TIMEs (**Fig.5c**) using Linear amplitude neuromodulation^27, 41^, Sinusoidal pulse-width neuromodulation^10, 91^, Poisson frequency neuromodulation (i.e. Poisson spiking train with mean frequency of 50 Hz, consisting in a non-biomimetic, frequency-variant stimulation, where spikes intervals are uncorrelated and exponentially distributed) and Biomimetic neurostimulation patterns constructed using FootSim (SAI-like, SAII-like, FAI-like, FAII-like and FULL Biomimetic).

The stimulation was delivered from 3 ASs for S1 and S2 eliciting sensation in the Frontal met, 3 AS for S1 and S2 eliciting sensation in the Central met, 3 AS for S1 and 2 AS S2 eliciting sensation in the Lateral met and 5 AS for S1 and 2 ASs S2 eliciting sensation in the Heel. For S3, only one AS per the four areas were tested (**Fig.S3**). The subjects were asked to report the location (i.e., Projected Field) and naturalness, rated on a scale from 0 to 5^27, 41, 56^. Each condition was randomized, and each stimulation trial was repeated three times. The injected charge (amplitude and pulse-width) was specific for each channel and set to the related threshold charge^36^. Moreover, intensity ratings were also collected during each stimulation to exclude relevant intensity difference among the encoding strategies (intensity bias). For the typical time scales involved in our experiments (trials lasting on the order of minutes), neither of our participants reported relevant changes in sensation intensity, which would indicate the presence of adaptation. The specific quality descriptors of the electrically-evoked sensations reported by the subjects were electrode-dependent, including a multitude of sensation types (natural and unnatural)^36, 92^. The subjects were blinded to the sensory encodings used in each trial.

### Real-time biomimetic neurostimulation in a neuro-robotic leg

The neuroprosthetic system included a robotic leg with a sensorized insole with embedded pressure sensors, along with a microcontroller and a neural stimulator^88^, implementing the encoding strategies and providing sensory feedback in real time by means of implanted TIMEs^36^. We implemented and tested: (i) no feedback (NF): the prosthesis did not provide any sensory feedback; (ii) linear amplitude neuromodulation (LIN): the prosthesis provided a linear feedback from three channels of the sensorized insole (heel, lateral or medial and frontal; more details in Petrini et al.^36^); iii) time-discrete neuromodulation feedback (DISC): the prosthesis delivered short trains of stimulation (0.5s) when a specific sensor was activated (heel, lateral or central and frontal) and again (0.5s) when the load was released from that sensor (neurostimulation delivered only at the transients); iv) biomimetic neuromodulation feedback (BIOM): the neuroprosthetic device provided the biomimetic stimulation, reported as the one eliciting more natural sensation, from three channels of the sensorized insole (heel, lateral or central and frontal). For the model-based biomimetic approach (BIOM), the corresponding frequency trains were computed previously offline by the model to reach the appropriate speed during the real-time implementation. The amplitude of the stimulation was modulated linearly with the pressure sensor output, as proposed in Valle et al., (HNM-1)^41^. In LIN and DISC, the stimulation frequency was fixed (tonic stimulation) to 50 Hz^7^. During the functional experiments reported in this work, three tactile channels (those eliciting sensation on the heel, lateral or medial and frontal met areas) were used for sensory feedback in all the conditions. The delivered charge was similarly modulated on the three stimulating channels, but in a different range. In fact, each channel was modulated between its threshold and maximum charge values identified in the last mapping session. The biomimetic stimulation patterns adopted on the three channels were selected according to the naturalness perceived per foot area (**Fig.S3**) in each implanted subject. In particular, FAI Biomimetic for frontal, lateral and heel for both S1 and S2, while FULL Biomimetic neurostimulation for lateral met in both S1 and S2.

### Stairs Task

During the stairs test (ST), S1 and S2 were asked to go through a course of stairs in sessions of 30s per 10 times per condition. The setup was configured as an angular staircase endowed with six steps with a height of 10 cm and a depth of 28 cm on one side and with four steps with a height of 15 cm and a depth of 27.5 cm on the other. Subjects were asked to walk clockwise climbing up the six steps and going down the four steps (**Fig.6a**). Walking sessions were performed in four distinct conditions: (i) no feedback (NF); (ii) linear neuromodulation feedback (LIN); iii) time-discrete neuromodulation feedback (DISC); iv) biomimetic neuromodulation feedback (BIOM). All the stimulation conditions were randomly presented to the volunteers. The gait speed for this task was reported in terms of number of laps, as previously performed^36, 49^. A lap is intended as going up and down the stairs and reaching the starting position again. A higher number of completed laps is indicative of a higher speed and vice versa. S1 and S2 performed this task.

### Cognitive double task

In the cognitive double task (CDT), first S1 and S2 were instructed to walk forward for 5 m (Baseline, **Fig.S4**) while timing them for 10 times per 4 conditions (BIOM, LIN, DISC and NF) performed in a random order. Subsequently, they were asked to walk for the same distance while performing a dual task (CDT). In particular, they had to spell backward in their mother-tongue language (Serbian) a five-letter word, which had not been previously presented. Also, this task was performed 10 times per 4 conditions (BIOM, LIN, DISC and NF) performed in a random order. While the subjects were performing the CDT, both the walking speed (m/s) and the accuracy of the spelling (% of correct letters) were recorded (**Fig.6b**). S1 and S2 performed this task.

### Self-reported confidence

At the end of each session of ST, participants were asked to assess their self-confidence while performing the motor task, using a visual analog scale (from 0 to 10). The data were acquired in BIOM, LIN, DISC and NF conditions in S1 and S2.

### Statistics

All data were exported and processed offline in Python (3.7.3, the Python Software Foundation) and MATLAB (R2020a, The MathWorks, Natick, USA). All data were reported as mean values ± SD (unless elsewise indicated). The normality of data distributions was verified. In case of Gaussian distribution, two-tailed analysis of variance (ANOVA) test was applied. Elsewise, we performed the Wilcoxon rank-sum test. Post-hoc correction was executed in case of multiple groups of data. Significance levels were 0.05 unless differently reported in the figures’ captions. In the captions of the figures, we reported the used statistical tests for each analysis and its result, along with the number of repetitions (n) and p values for each experiment.

## Acknowledgments

The authors are deeply grateful to the three subjects who freely donated months of their life for the advancement of knowledge and for a better future for leg amputees. We thank Prof. Marco Capogrosso for the support during the animal experimentation and the related data analysis. We also thank the National Centre of Competence in Research (NCCR) Robotics for the useful collaborations. The funder had no role in the experimental design, analysis, or manuscript preparation or submission. All authors had complete access to data. All authors authorized submission of the manuscript, but the final submission decision was made by the corresponding authors. This project has received funding from the European Research Council (ERC) under the European Union’s Horizon 2020 research and innovation program (FeelAgain grant agreement no. 759998), from Gebert Ruf Stiftung (InnoBooster, MYLEG, GRS–096/21), from Swiss National Science Foundation (SNSF) (MOVE-IT no. 197271), from project IDEJE by Science Fund of the Republic of Serbia (DiabeticReTrust no. 7753949), by project (ID: 93022925/ 94030803) of the St. Petersburg State University, St. Petersburg (for N.P.), by Sirius University of Science and Technology project: NRB-RND-2115 (for P.M.). The work was carried out within the framework of the Implementation Program Priority 2030 (NUST MISIS) (for O.G.).

## Author contributions

G.V. developed the neurostimulation software for human and animal experiments, developed the biomimetic neuroprosthetic leg systems, performed the human experiments, analyzed the data collected in humans, supervised all the analyses, prepared the figures and wrote the paper; N.K. developed the in-silico model (FootSim), generated the tested stimulation patterns using FootSim, supervised and performed the analyses on the animal data, prepared the figures and wrote the paper; D.E. performed the analyses on the animal data and prepared results related the figures; N.P. performed the animal surgical procedures. O.G. performed the animal experiments; P.C. and T.S. developed the TIME and delivered technical assistance for the human implantation and explanation procedures and reviewed the manuscript; P.M. designed and performed the animal surgical procedures and performed the animal experiments; M.B. performed the human surgeries and was responsible for all the clinical aspects of the human study; S.R. designed the study, performed and supervised the human and animal experiments, supervised the analyses, wrote the paper. All authors edited and proofread the manuscript.

## Data and materials availability

The authors declare that the data supporting the findings of this study are available within the article and its supplementary information files. Software routines developed for the analysis are available from the corresponding author. Other data can be made available to qualified individuals for collaboration provided that a written agreement is signed in advance between the included consortium and the requester’s affiliated institution.

