## Supplementary material for "Biomimetic computer-to-brain communication restoring naturalistic touch sensations via peripheral nerve stimulation"

† Equal contribution

**List of Supplementary Materials:**

Fig. S1. Synchronization of the neural activity evoked in the spinal cord.

Fig. S2. Biomimetic neurostimulation elicits more natural sensations than non-biomimetic approaches in Subject 3.

Fig. S3. Electrically-evoked sensation naturalness related to different projected fields location.

Fig. S4. Walking speed baseline in the Cognitive Dual Task (CDT).

Table S1. Participants’ demographics.

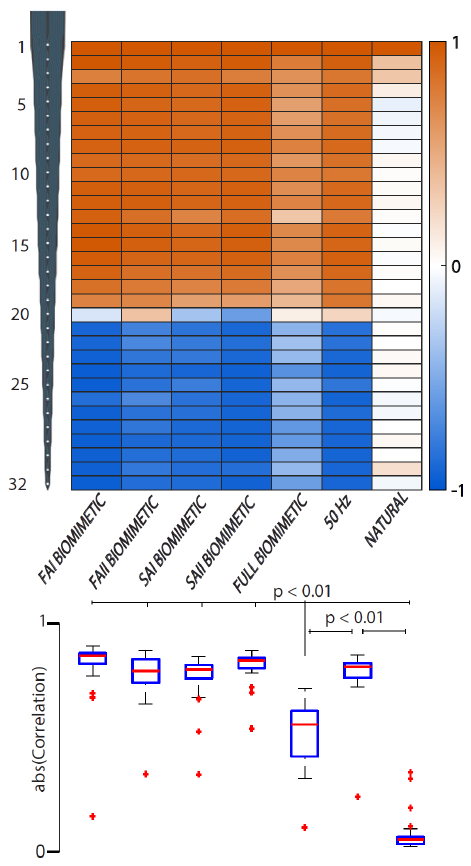

**Fig S1. Synchronization of the neural activity evoked in the spinal cord.** We compared and presented the correlation between the neural local field potential recording in the first electrode channel with all the other channel recordings. Upper part: Correlation values are color coded and presented along the spinal array electrode axes. Bottom part: We compared absolute values of correlation coefficient of natural with every biomimetic condition as well as 50 Hz stimulation with FULL biomimetic and natural touch condition with one-way ANOVA. Boxplots: The central mark indicates the median, and the bottom and top edges of the box indicate the 25th and 75th percentiles, respectively. The whiskers extend to the most extreme data points not considered outliers, and the outliers are plotted individually using the red '+' symbol. p values are indicated

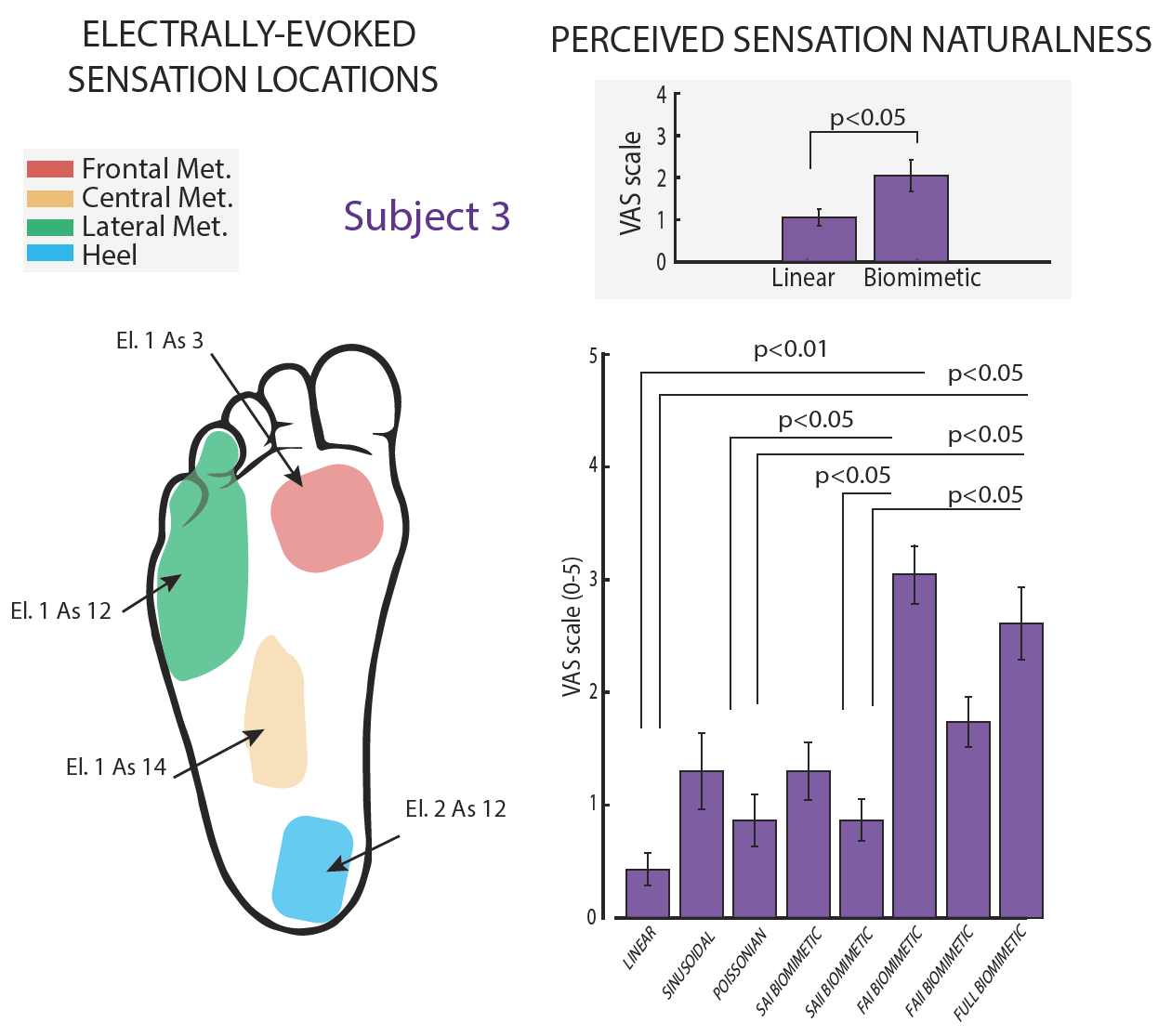

**Fig S2. Biomimetic neurostimulation elicits more natural sensations than non-biomimetic approaches in Subject 3.** Projective fields map of Subject 3 related to the active sites adopted to electrically stimulate the tibial nerve. Different colors show the 4 main regions of the phantom foot (Frontal, Lateral and Central Metatarsus, and Heel). Naturalness ratings (VAS scale 0-5) of the perceived sensation elicited exploiting different stimulation strategies. Insets: Group comparison between linear vs biomimetic stimulations.

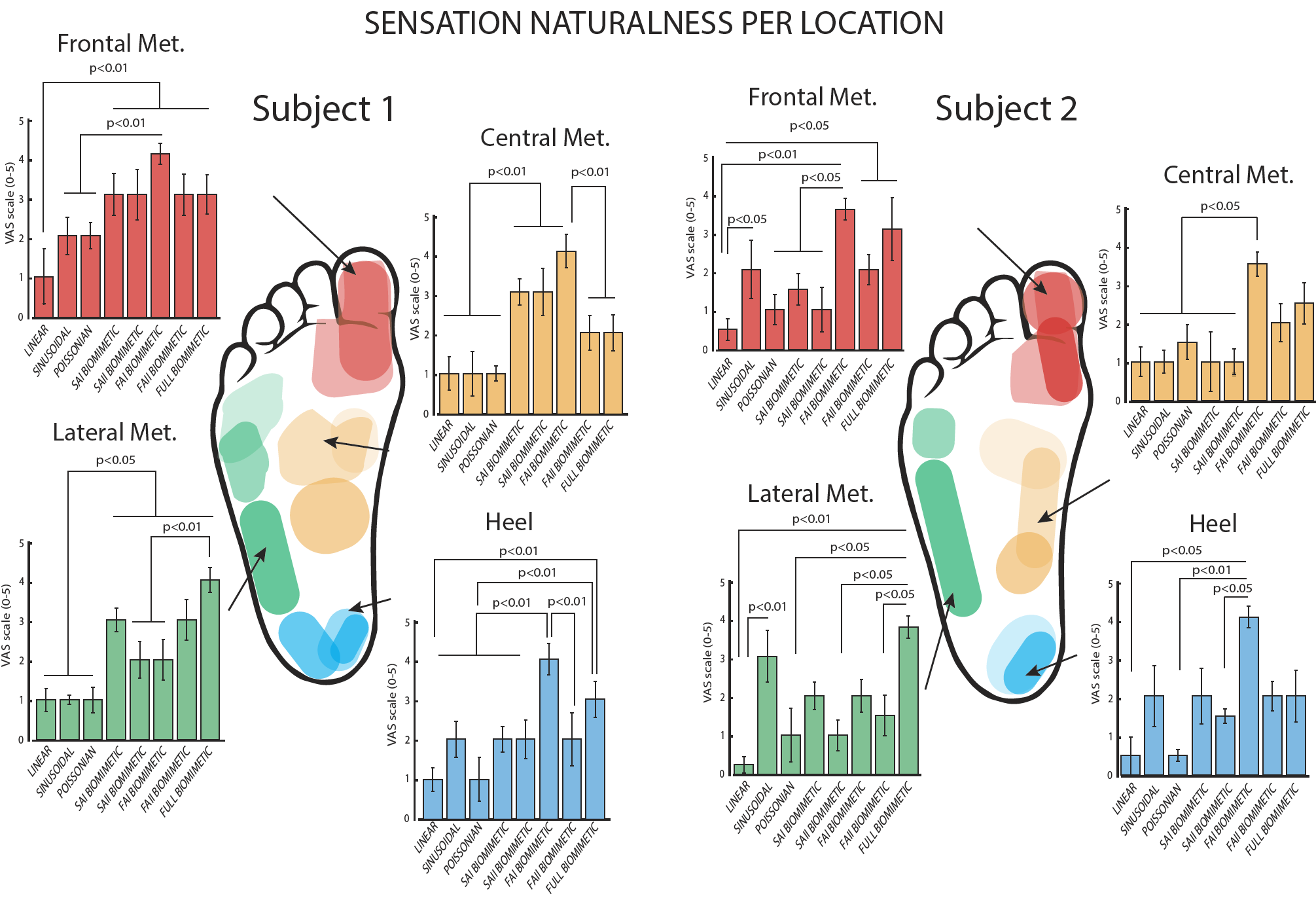

**Fig S3. Electrically-evoked sensation naturalness related to different projected fields location.** Projective fields maps of two implanted subjects (1 & 2) related to the active sites adopted to electrically stimulate the nerves. Different colors show the 4 main regions of the phantom foot (Frontal, Lateral and Central Metatarsus, and Heel). Naturalness ratings (VAS scale 0-5) of the perceived sensations breakdown per location of the projected fields.

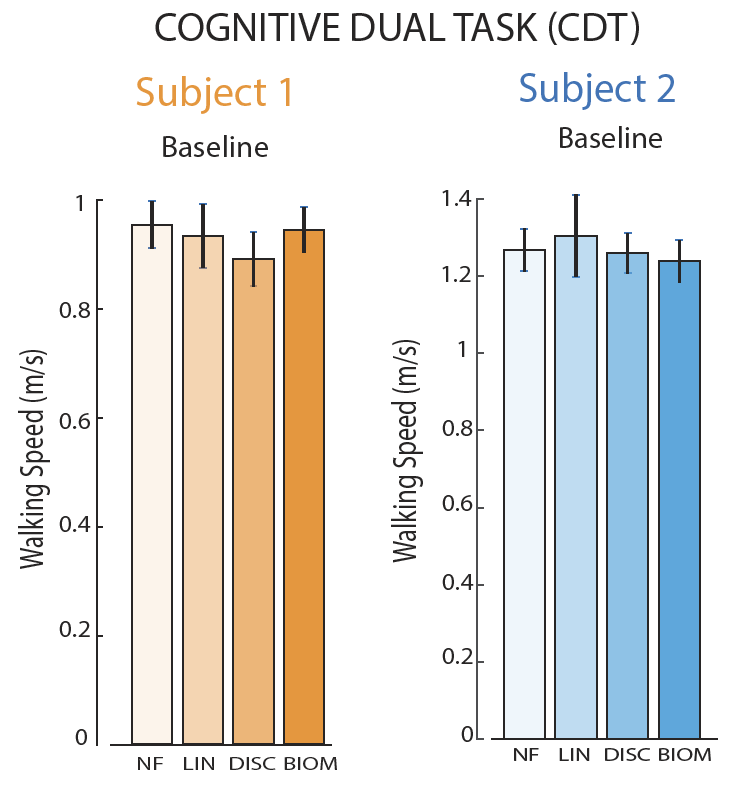

**Fig S4. Walking speed baseline in the Cognitive Dual Task (CDT).** Motor performance (Walking Speed – m/s) of Subject 1 & 2 in the Cognitive Dual Task (CDT) at baseline (no mental task). The tested conditions are NF (No Feedback), LIN (Linear Neurostimulation), DISC (Discrete Neurostimulation) and BIOM (Biomimetic Neurostimulation).

**Table S1.** **Participants’ demographics.**

| Patient | Cause of  Amputation | Level and Side of Amputation | Age | Amputation Time | Phantom Limb Pain | Gender | Own Prosthesis | Frequency of Use |
| --- | --- | --- | --- | --- | --- | --- | --- | --- |
| S1 | Trauma | Distal two-thirds of right thigh | 49 | 2 | Medium | M | Passive prosthesis (3R80-Ottobock) | Daily |
| S2 | Trauma | Distal two-thirds of right thigh | 35 | 12 | Medium | M | Passive prosthesis (3R80-Ottobock) | Daily |
| S3 | Trauma | Distal two-thirds of left thigh | 53 | 7 | Low | M | Passive prosthesis (3R80-Ottobock) | Daily |
